# bletl - A Python Package for Integrating Microbioreactors in the Design-Build-Test-Learn Cycle

**DOI:** 10.1101/2021.08.24.457462

**Authors:** Michael Osthege, Niklas Tenhaef, Rebecca Zyla, Carolin Müller, Johannes Hemmerich, Wolfgang Wiechert, Stephan Noack, Marco Oldiges

## Abstract

Microbioreactor (MBR) devices have emerged as powerful cultivation tools for tasks of microbial phenotyping and bioprocess characterization and provide a wealth of online process data in a highly parallelized manner. Such datasets are difficult to interpret in short time by manual workflows. In this study, we present the Python package bletl and show how it enables robust data analyses and the application of machine learning techniques without tedious data parsing and preprocessing. `bletl` reads raw result files from BioLector I, II and Pro devices to make all the contained information available to Python-based data analysis workflows. Together with standard tooling from the Python scientific computing ecosystem, interactive visualizations and spline-based derivative calculations can be performed. Additionally, we present a new method for unbiased quantification of time-variable specific growth rate 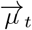 based on a novel method of unsupervised switchpoint detection with Student-t distributed random walks. With an adequate calibration model, this method enables practitioners to quantify time-variable growth rate with Bayesian uncertainty quantification and automatically detect switch-points that indicate relevant metabolic changes. Finally, we show how time series feature extraction enables the application of machine learning methods to MBR data, resulting in unsupervised phenotype characterization. As an example, t-distributed Stochastic Neighbor Embedding (t-SNE) is performed to visualize datasets comprising a variety of growth/DO/pH phenotypes.

**Practical Application:** The `bletl` package can be used to analyze microbioreactor datasets in both data analysis and autonomous experimentation workflows. Using the example of BioLector datasets, we show that loading such datasets into commonly used data structures with one line of Python code is a significant improvement over spreadsheet or hand-crafted scripting approaches. On top of established standard data structures, practitioners may continue with their favorite data analysis routines, or make use of the additional analysis functions that we specifically tailored to the analysis of microbioreactor time series.

Particularly our function to fit cross-validated smoothing splines can be used for *on-line* signals from any microbioreactor system and has the potential to improve robustness and objectivity of many data analyses. Likewise, our random walk based 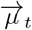 method for inferring growth rates under uncertainty, but also the time-series feature extraction may be applied to *on-line* data from other cultivation systems as well.

## 1 Introduction

The development of innovative bioprocesses is nowadays often carried out in a Design – Build – Test – Learn (DBTL) cycle [1], where fast iterations of this cycle are desired to shorten development times and therefore safe costs. This acceleration can be enabled by modern genetic engineering tools, lab automation and standardized data analysis pipelines. One aspect in the “Test” part of the DBTL cycle of a bioprocess is the cultivation of the microorganisms to be tested. This is often performed in microbioreactor systems, since they provide a good balance between adequate throughput and scalability of the results to laboratory scale bioreactors as the gold standard [2][3].

A typical microbioreactor system will provide transient monitoring of biomass formation, dissolved oxygen, pH, and fluorescence. Usually, the researcher has access to additional environmental data such as temperature, shaking or stirrer frequencies, humidity, and gas atmosphere. Analyzing this heterogeneous, multi-dimensional data in a quick and thorough manner can be challenging, especially since vendor software often covers only a limited amount of use cases.

From our experience, most researchers try to alleviate such problems by employing more-or-less sophisticated spreadsheets, available with various software solutions. While presenting an easy way for simple calculations and visualizations, extensive analysis of the data quickly results in hardly maintainable documents, which are impossible for colleagues to comprehend, error-prone and easy to break. Most importantly, such multi-step manual data transformations do not comply with the FAIR data principles, whereas our `bletl` package directly addresses the accessibility aspect and creates incentives to, for example, retain the original data.

Automated data analysis pipelines solve this problem by removing the repetitive and error-prone manual workflows in favor of standardized workflows defined in code. Such workflows offer many advantages, if done correctly: a) data processing is clearly understandable and documented, b) every step is carried out for every input data file in the same way, guaranteeing the integrity and reproducibility of the results, c) data processing can be autonomously started after data generation and d) such a pipeline can be run on remote systems, which is especially useful for computational demanding calculations. Such data analysis pipelines are routinely used, *e*. *g*., for sequencing data [4], but seldom used for microbioreactor data. One example is the automated calculation of growth rates using MATLAB [5].

Additionally, automated data analysis opens up possibilities for at-line analysis and subsequent intervention during an experiment: Cruz Bournazou *et al*. report a framework for online experimental redesign using a data pipeline implemented in MATLAB [6]. Jansen *et al*. build a Python-based process control system to control pH and enzyme addition to a microbioreactor system using a liquid handling robot [7]. While those examples can automatically parse and analyze microbioreactor data, they are tailored to a specific use case and hence not universally applicable.

In this study, we introduce `bletl` as an open-source Python package for standalone and at-line parsing and analysis of microbioreactor data. The name `bletl` is inspired by the fact that it simplifies the implementation of <u>e</u>xtract, <u>t</u>ransform, <u>l</u>oad data processing workflows specifically for BioLector datasets. It is capable of parsing raw data without involving vendor software, making necessary calibrations for fluorescent-based measurement of pH and dissolved oxygen, and presenting all measurement, environmental and meta data in the easily accessible, `DataFrame` format from the popular `pandas` library [8, 9]. Currently, `bletl` is designed to parse data from the devices BioLector I, II, and Pro manufactured by Beckman Coulter Life Sciences, but its general design and methods can be applied also for other devices. Its analysis submodules build on top of `bletl`’s standardized data structures and provide the user with sophisticated methods for microbioreactor data analysis, such as spline approximation with cross validation for data smoothing, growth rate analysis using Bayesian modeling techniques as well as time series feature extraction. In addition to an extensive documentation and automated software tests, we provide application examples using relevant experimental data. With `bletl`, scientists using microbioreactors have a powerful tool to make their data analysis less cumbersome and error-prone, while they can directly benefit from state-of-the-art machine learning techniques.

## 2 Materials and Methods

### 2.1 Core package

The `bletl` package includes the data structures that are common to datasets originating from BioLector I, II and Pro microbioreactors. Parsing of raw data is deferred to *parsers* that may implement logic that is specific to a certain BioLector model or file type.

#### 2.1.1 Parsing and data structures

Parsing of raw data typically begins with a call to the `bletl.parse` function, which first determines the file type from its content. The parsing procedure does not only ingest the data into accessible Python data structures, but also takes care of re-naming tabular data columns to a standardized naming scheme and type-casting values to integer, float or string types. After the BioLector model and file type version are identified, a matching *parser* is selected, thereby enabling a plug-in system for specialized parsers for different file type versions or the new BioLector XT device. The parsing logic is highly dependent on the BioLector model, but generally follows the pattern of first separating the file header of metadata from the table of measurement data. Logical blocks of information, such as the table of filtersets are then parsed into `pandas.DataFrame` objects. These tabular data structures are collected as attributes on a `bletl.BLData` object which is returned to the user. The `BLData` class is a Python dictionary data type with additional `properties` and `methods`. Via its properties, the user may access various `DataFrame` tables of relevant metadata, including the aforementioned tables of filtersets, comments or environment parameters such as chamber temperature or humidity.

Key-value pairs in the `BLData` dictionary are the names of filtersets and corresponding `FilterTimeSeries` objects. This second important type from the `bletl` package hosts all measurements that were obtained with the same filterset. Like the `BLData` class it provides additional methods such as `FilterTimeSeries.get_timeseries` for easy access of time/value vectors.

### 2.2 Analysis methods

In submodules of the `bletl` package, various functions are provided to facilitate higher-throughput and automated data analysis from bioprocess timeseries data.

One often used feature is the `find_do_peak` function that implements a Dissolved Oxygen (DO)-peak detection heuristic similar to the one found in the m2p-labs *RoboLector* software. The DO-peak detection algorithm finds a cycle number corresponding to a DO rise, constrained by user-provided threshold and delay parameters.

Additional, more elaborate analysis functions were implemented to allow for advanced data analysis or experimental control.

#### 2.2.1 Spline approximations

To accommodate for the measurement noise in *on-line* measured time series, various smoothing procedures may be applied to the raw signal. A popular choice for interpolation are spline functions, specifically *smoothing splines* that can reproduce reasonable interpolations without strong assumptions about the underlying relationship. With `bletl.get_crossvalidated_smoothing_spline` we implemented a convenience function for fitting smoothing splines using either `scipy` or `csaps` [10, 11] for the underlying implementation. Both smoothing spline implementations require a hyperparameter that influences the amount of smoothing. Because the choice of the smoothing hyperparameter strongly influences the final result we automatically apply stratified k-fold cross-validation for determining its optimal value. The implementation can be found in the code repository of the `bletl` project [12].

#### 2.2.2 Growth rate analysis

A “calibration-free” approach to calculate time-variable specific growth rate *μ*(*t*) (1) relies on the previously introduced spline approximations, combined with the popular assumption of a linear backscatter *Y_BS_* vs. biomass *X* relationship (2).

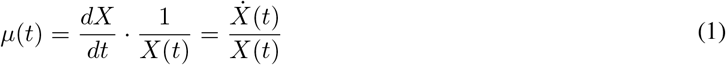

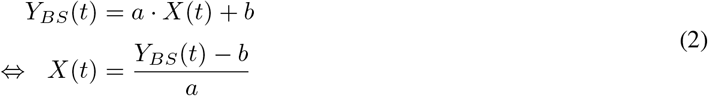

Substituting the biomass *X*(*t*) in (1), the slope parameter *a* cancels out such that only a “blank” *b* and the measured backscatter *Y_BS_*(*t*) are needed for a specific growth rate calculation (3).

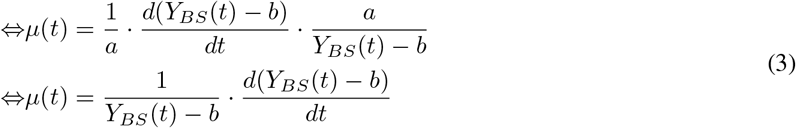

Finally, the backscatter curve *Y_BS_*(*t*) can be approximated by a smoothing spline *S*_YS,blanked_ (*t*) to obtain a differentiable function (4).

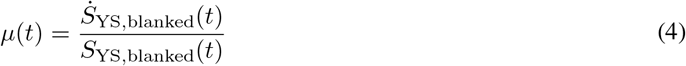

An alternative approach is to construct a *generative* model of the biomass growth. In essence, the time series of observations is modeled as a deterministic function of an initial biomass concentration *X*_0_ and a vector of specific growth rates 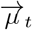 at all time points where observations were made. The structure of this model assumes exponential growth between the time steps, which is a robust assumption for high-frequency time series such as the ones obtained from BioLector processes.

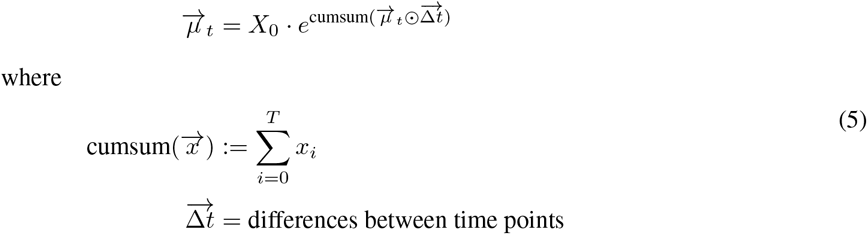

The convenience function `bletl.growth.fit_mu_t` creates the generative 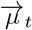 model (5) from user-provided vectors of observation time points 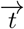 and backscatter values 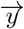. To contrast it from the smooth, continuous *μ*(*t*) from (4) we use the 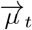 notation to underline that the approach discretizes the growth rate into a step function. The model is built with the probabilistic programming language PyMC3 [13, 14] and an optimal parameter set, the maximum *a-posteriori* (MAP) estimate, is found automatically. Additionally, the user may decide to perform Markov-chain Monte Carlo (MCMC) sampling using advanced sampling algorithms such as No-U-Turn Sampler (NUTS) from the PyMC3 [13] package to infer probability distributions for the model parameters *X*_0_ and 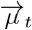.

In the generative *μ_t_* model, the vector of growth rates is modeled with either a Gaussian or Student-t distributed random walk (Figure 1). This does not only result in a smoothing of the growth rate vector, but enables additional flexibility with respect to switchpoints in the growth rate. A `drift_scale` parameter must be given to configure the random walk with a realistic assumption of how much growth rate varies over time. Small `drift_scale` corresponds to the assumption that growth rate is rather stable, whereas large `drift_scale` allows the model to describe a more fluctuating growth rate distribution. Additionally, the user may provide previously known time points at which growth rate switches are expected. Examples of such switchpoints are the time of induction, occurrence of oxygen limitation or the time at which the carbon source is depleted. If 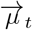 is described by a Student-t random walk, switchpoints can be detected automatically by inspecting the prior probabilities of the estimated growth rate in every segment (Figure 1). Our implementation automatically classifies elements of 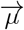 as switchpoints as soon as their prior probability is < 1 %.

**Figure 1:**
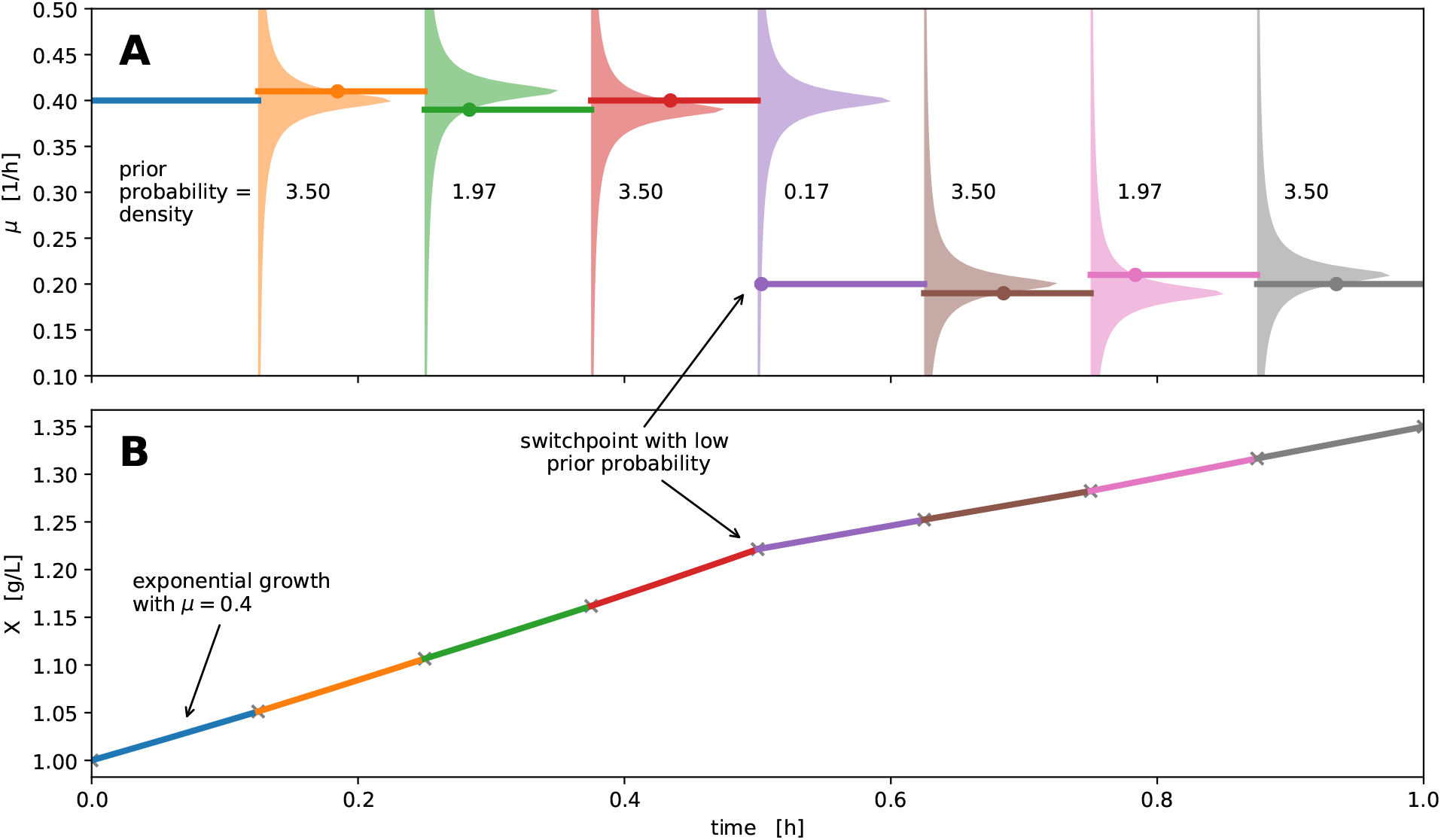
Switchpoint detection with a Student-t random walk. A microbial growth curve with fluctuating specific growth rate *μ* may be discretized into segments of exponential growth with constant growth rate (solid lines). Modeling such a sequence of growth rates with a random walk assigns prior probabilities to every value of 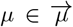, centered on the values of previous iterations (**A**, colored areas). When fitting the model these prior probabilities “pull” subsequent values in 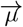 towards each other, leading to a smoothing and counteracting overfitting. Using a fat-tailed Student-t distribution for the random walk prior, the penalty for large jumps is less extreme compared to a Normal distribution, thereby allowing for jumps in the random walk (5th segment).

While there has been prior work on using random walks for outlier detection [15] we are not aware of any prior work using Student-t random walks for the unsupervised detection of switchpoints in time series data.

#### 2.2.3 Feature extraction

The `bletl.features` submodule implements functions for the automated extraction of both biologically and statistically motivated time series features. An abstract `Extractor` class may be inherited to implement feature extraction of characteristic features such as DO peaks. Additionally, our `TSFreshExtractor` uses the open source Python package `tsfresh` [16] to extract hundreds of features from variable length time series automatically. Starting from a `bletl.BLData` object containing one or more `FilterTimeSeries`, the `bletl.features.from_bldata` function extracts features from multiple filtersets using a user-specified mapping of `Extractor` objects. Optionally, a dictionary of well-wise cycle numbers can be passed to truncate time series to the relevant cycles. The results are returned as a `DataFrame` for maximal compatibility with downstream analysis operations. For details on the implementation we refer to the code and documentation [12, 17].

Figure 2 shows how the function is applied to our demonstration data set. The resulting `DataFrame` comprised 2343 feature columns for each of the 48 wells in the input data. Feature columns with `NaN`, ±∞ entries or without variability in their values were dropped, resulting in 1282 features available for further analysis.

**Figure 2:**
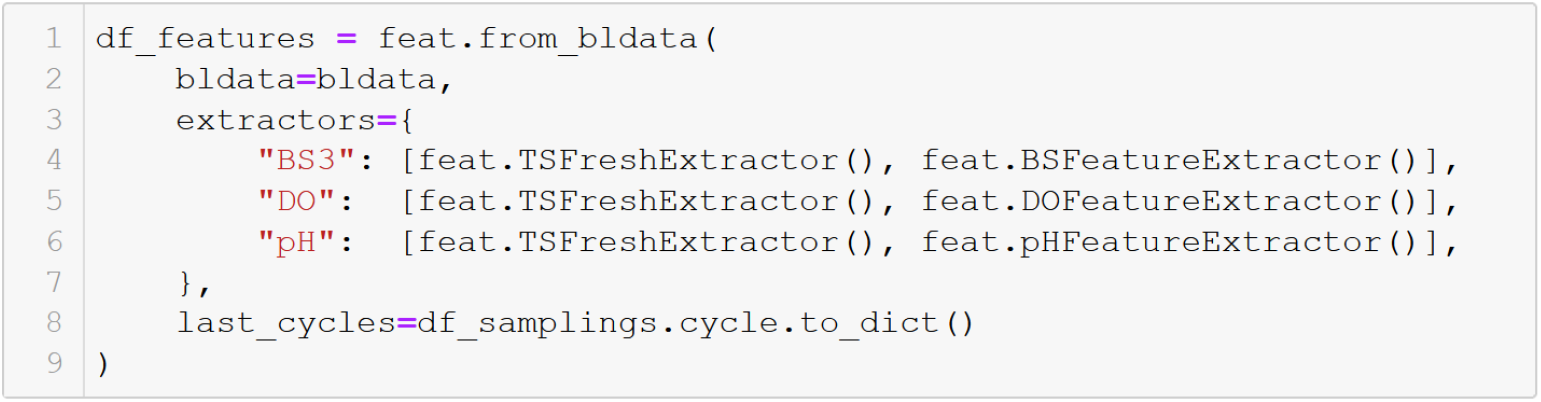
Code to run feature extraction from three filtersets. The name of each filterset is mapped to a list of `Extractors` that may include user-defined feature extraction implementations. The `last_cycle` keyword argument can be used to pass a mapping of well IDs to the last relevant cycle numbers. Extracted features are returned in the form of a `pandas.DataFrame`.

#### 2.2.4 Visualization by t-SNE

Starting from features extracted with `bletl.features` we applied the t-SNE technique to find a two-dimensional embedding for the visualization of local structure in the dataset. The extracted time series features were cleaned such that features with `NaN` values, or without diversity were removed. After feature-cleaning, the t-SNE implementation from scikit-learn was applied with a perplexity setting of 10 and initialization by PCA.

### 2.3 Media and cultivation conditions

The dataset presented as an application example in this study was obtained in an automated cultivation workflow on the previously described microbial phenotyping platform. 48 cultures of *Corynebacterium glutamicum* ATCC 13032 harboring the pPBEx2[18]-based plasmid pCMEx8-NprE-Cutinase (sequence in section 4.3) were cultivated in CGXII medium (recipe as in [19]) with different carbon sources. Carbon sources were prepared as C-equimolar, random combinations of 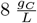 glucose, fructose, maltose, sucrose, gluconate, lactate, glutamate or *myo*-inositol. For every well except A01, where glucose was the sole carbon source, three different carbon sources were chosen at random. A total of 140 μL of C-equimolar carbon source stock were added to each well. The 140 μL were split into 7 parts of 20 μL and such that at least one part was used for each selected carbon source and the remaining 4 parts were assigned randomly. The resulting media composition in terms of pipetted volume, and carbon mass per μL can be found in Section 4.1.

800 μL CGXII medium were inoculated to an optical density at 600 nm of 0.2. Cultures were grown in a MTP-48-BOH 1 FlowerPlate in a BioLector Pro (both Beckman Coulter Life Sciences, USA) at 1400 rpm, 30 °C and ≥ 85 % humidity. Expression was induced autonomously with 10 μL Isopropyl-*β*-D-thiogalactopyranosid (IPTG) (final concentration 100 μM) when cultures reached a backscatter value of 5.82, corresponding to approximately 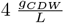. Culture from each well was harvested 4 hours after induction [20].

## 3 Results and Discussion

### 3.1 Basic visualization workflow

Every analysis begins with loading data into a structure that can be used for further analysis. In the case of a BioLector experiment, the data are multiple tables that hold information about filtersets, environment variables such as temperature or humidity, as well as the well-wise measurements. In most cases the result files already contain relevant meta information such as lot number or process temperature and parsing them with `bletl` comes down to a single line of Python (Figure 3).

**Figure 3:**
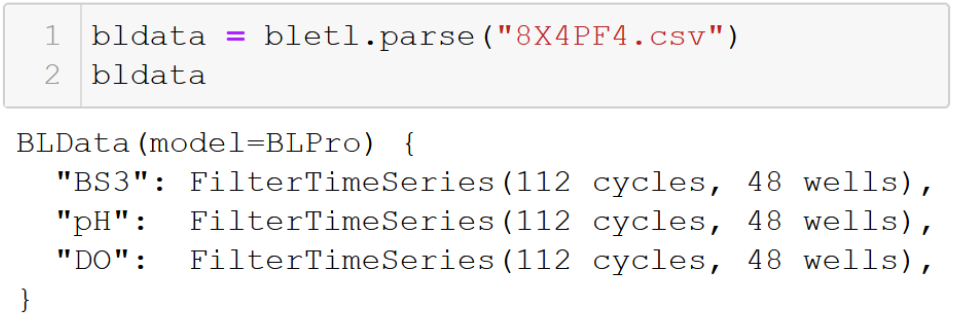
Parsing of a BioLector result file. The `bletl.parse` function automatically determines the file type (BioLector I, II or Pro) and applies calibration of optode measurements based on lot number and temperature from the file. Optionally, lot number and temperature, or calibration parameters may be passed to override the values from the file. The function can also process a list of result file paths and automatically concatenate them to a single `BLData` object.

The `bletl.BLData` type is a dictionary-like data structure into which results are loaded. It has additional properties through which process metadata and measurements that are not tied to individual wells can be obtained. Its text representation, which appears when the object is displayed in an interactive Jupyter notebook session shows the names of filtersets and the amount of data they contain (Figure 3).

Elements in the `BLData` object are the `FilterTimeSeries`, that contain the well-wise measurements. The simplest way to access the time series of a particular filterset and well is via the `BLData.get_timeseries(well, filterset)` or `FilterTimeSeries.get_timeseries(well)` methods. Optionally, a `last_cycle` number may be passed to retrieve only the data up to that specific cycle number. This is useful in situations where wells were sampled and might only be analyzed up to the sampling time point.

With the data structures provided by our `bletl` package, the data analysis workflow for a BioLector experiment is no different to any standard data analysis performed with Python. Figure 4 shows an example how a simple figure of process variables can be prepared using the popular `matplotlib` visualization library. Such a visualization of measurements, together with annotations of, for example, measurement pausing events can help to diagnose irregularities, or to provide a quick overview about the experiment.

**Figure 4:**
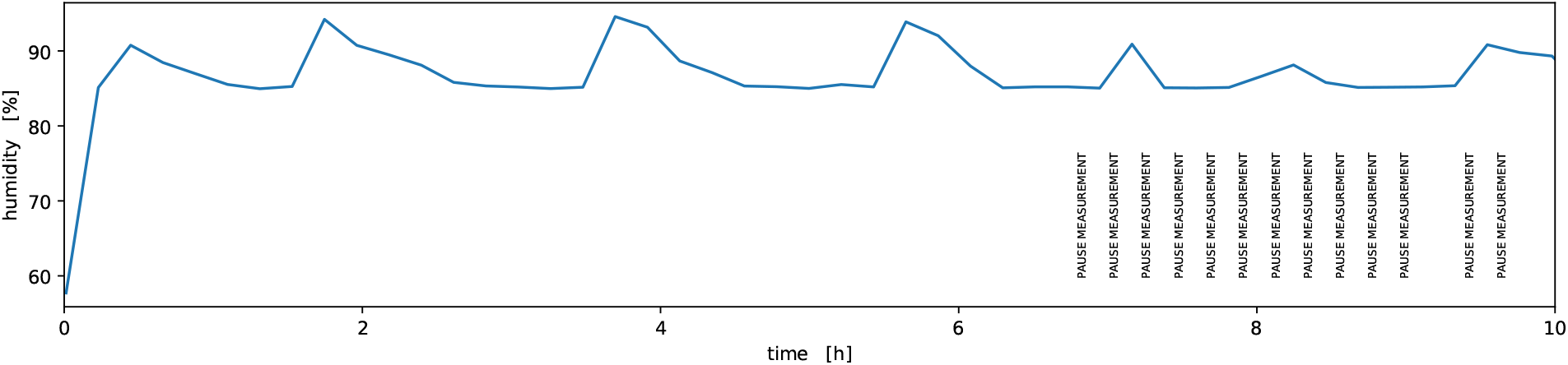
Visualization of process variables. This screenshot shows how a simple figure of humidity over time can be prepared alongside annotations of “pause measurement” events. In lines 4 and 5 the time and value vectors for humidity measurements are fetched from the `environment pandas.DataFrame` property of the previously loaded data object. The `for`-loop in lines 8-13 iterates over entries of another table `comments` that holds user- and system-comments of the BioLector process.

In the dataset presented here, cultures were induced and sampled by a robotic liquid handler. Induction was triggered *at-line* based on latest backscatter observations and sampling was performed 4 hours after the induction events Section 2.3. The meta data of these induction and sampling events were logged into an XLSX file and loaded into a `pandas.DataFrame` for the analysis Table 1.

**Table 1:** Excerpt of sampling event log

| well | timestamp | time | cycle | volume | supernatant_well |
| --- | --- | --- | --- | --- | --- |
| A01 | 2020-07-21T08:10:04.721Z | 13.076466 | 61 | -950 | H01 |
| A02 | 2020-07-21T07:05:04.299Z | 11.993016 | 56 | -950 | G01 |
| A03 | 2020-07-21T08:10:04.786Z | 13.076485 | 61 | -950 | F01 |
| A04 | 2020-07-21T08:10:04.841Z | 13.076500 | 61 | -950 | E01 |
| A05 | 2020-07-21T06:26:17.384Z | 11.346651 | 53 | -950 | D01 |

The meta information about induction and sampling events is important for the analysis, because backscatter, pH and DO observations made after a well was *sacrifice*-sampled must be truncated before analysis or visualization.

Most data analyses are driven by interactive exploration of the data. With `bletl`, or rather with a Python-based data analysis workflow in general, this is facilitated by interactive plots using helper functions from, for example, the `ipywidgets` library. Figure 5 shows the code and resulting interactive plot of measurement results from a BioLector dataset. In comparison to Figure 4 the example shown in Figure 5 is marginally more involved, but again relies on standard tooling from the Python ecosystem. In this case the `ipywidgets` library is used to wrap a plotting function and create interactive input elements for selecting the filterset and wells to show.

**Figure 5:**
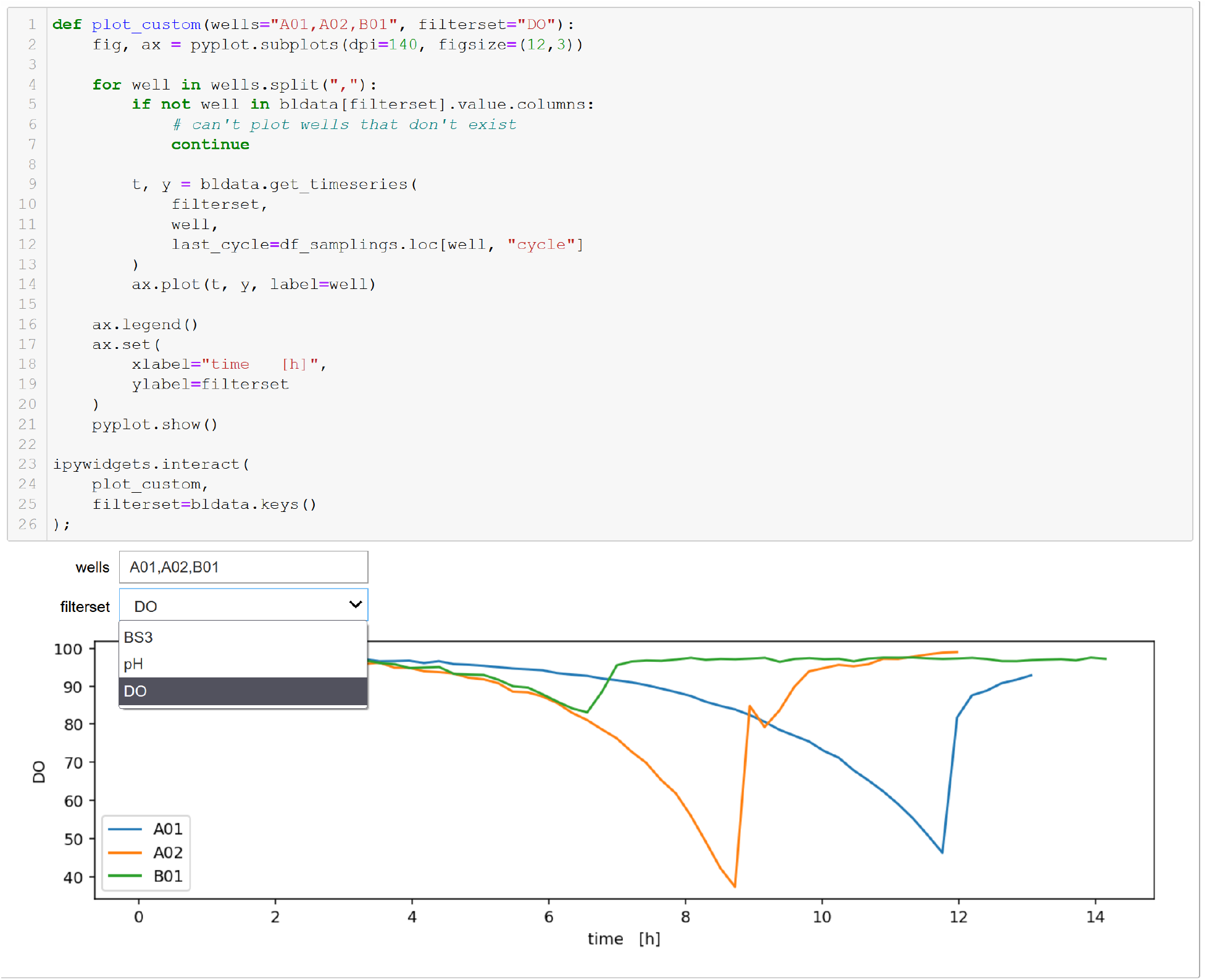
Interactive plot of well-wise measurements. A `plot_custom` function, defined in lines 1-17 takes a comma-separated text of well IDs and the name of a filterset as parameters for the visualization. In line 4 it iterates over the well IDs to create lines plots of the measurements, passing the number of the last relevant cycle from the event log (Table 1) to truncate the data. Line 21 passes the list of filtersets in the dataset (Figure 3) as options for the `fs` keyword-argument of the plotting function, thereby populating the dropdown menu.

### 3.2 Splines for time series smoothing and derivatives

Optical *on-line* measurements as those performed by the BioLector are inevitably subject to measurement noise. While the measurement noise of DO and pH signals in the BioLector II/Pro system was greatly reduced compared to the BioLector I model, it still requires special attention in subsequent data analysis procedures. Particularly in automated *at-line* decision making such as triggered induction or sampling, measurement noise can cause problems with threshold-based heuristics. With noisy *on-line* signals, such as optode measurements in BioLector I datasets, smoothing splines can yield more accessible visualizations and allow for finer-grained comparisons. Furthermore, the slope of the signals may be used for more sophisticated analysis or decisions.

For *at-line* triggers based on such noisy process values, a smoothing of the signal can increase the reproducibility of detecting, for example, a pH threshold. At the same time, the slope of process values often gives more process insight compared to absolute values alone. For example, a dissolved oxygen tension of 60 % alone is not very meaningful, but the observation of a strong positive slope tells the process engineer that the microbes might grow with reduced oxygen uptake rate. The calculation and visualization of pH and DO slopes is therefore an important tool for process data analysis.

Splines are a popular choice for both smoothing and derivative calculation, because they make few assumptions about the data and are available in most standard data analysis software. There are however multiple flavors of *smoothing splines* and they come with a smoothing parameter whose value has a considerable effect on the results. In `bletl.splines` we implemented a convenience function that automatically performs k-fold cross-validation on the smoothing parameter of either a `UnivariateSpline` cubic spline from `scipy` or a `UnivariateCubicSmoothingSpline` from *csaps* (cf. Section 2.2.1).

In Figure 6 the two spline methods were applied to pH and DO time series of well A01 from our demonstration dataset. Both smoothing spline methods find interpolations (solid lines) of the raw data that are almost indistinguishable. Their 1st derivative however reveals that the `UnivariateCubicSmoothingSpline` (ucss) from the csaps package is much more wiggly compared to the `UnivariateSpline` (us) from SciPy.

**Figure 6:**
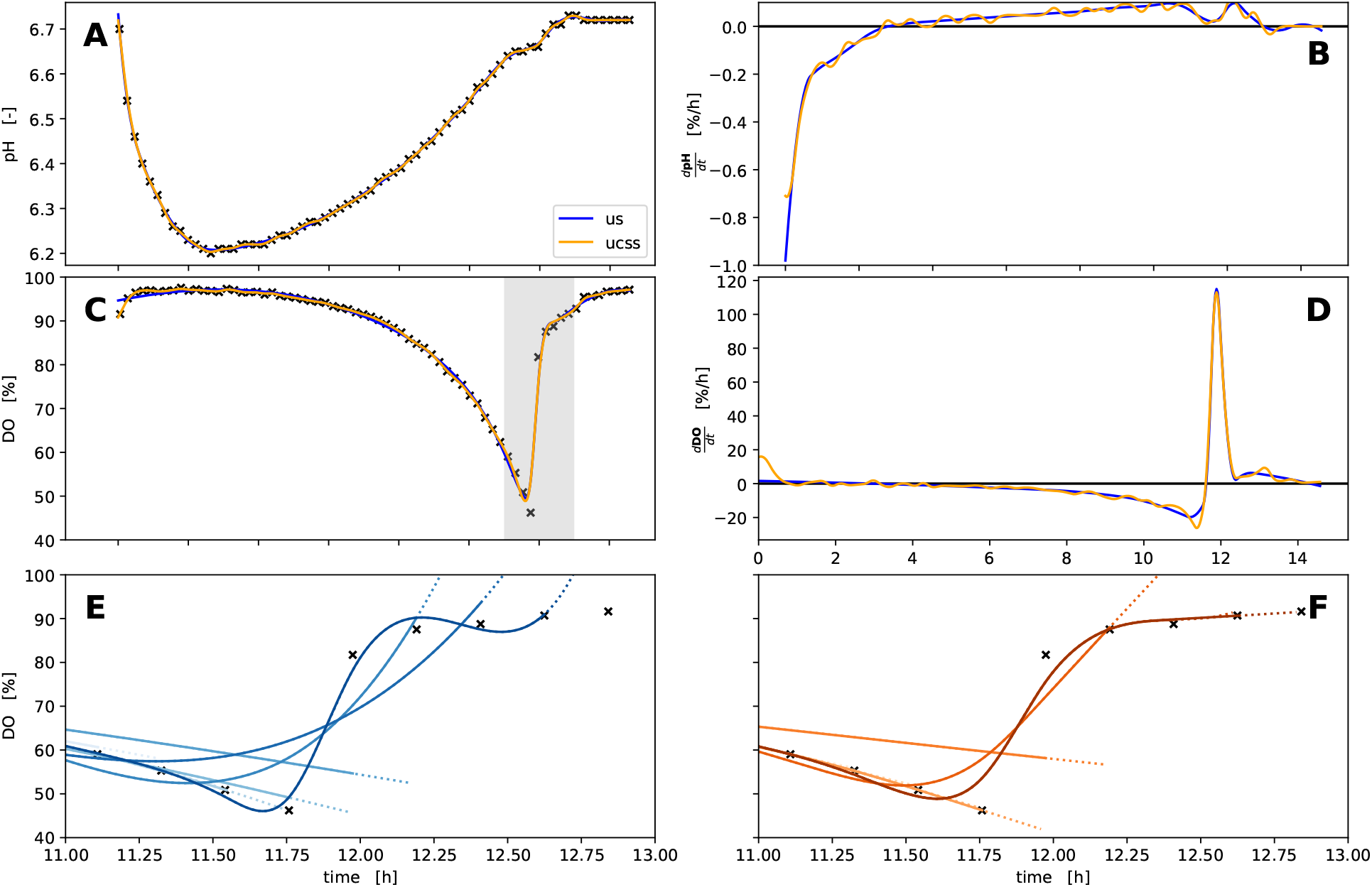
Splines fitted to pH and DO time series. Both spline methods `“us”` (blue) and `“ucss”` (orange) were applied to measurements of pH (**A**) and DO (**C**). The resulting reconstruction/interpolation is largely identical (**A, C**) with most notable differences between the methods at the start/end, as well as their derivatives (**B, D**), where the `“ucss”` method has considerably more *wiggly* derivatives. In the bottom row (**E, F**), the extrapolation (dotted lines) of splines fitted to data subsets of different length (solid lines) is straight for the `“ucss”` method, whereas the `“us”` method extrapolates with a curvature.

The bottom row shows a comparison of both methods in a simulated *at-line* situation where the DO time series grows point by point at the time of the characteristic substrate-depletion DO-peak. In this situation the `“us”` method produces strong alternating positive or negative slopes and curved extrapolation at the end of the curve. In contrast, the splines obtained with the `“ucss”` method extrapolate with an (almost) constant slope. Taking both scenarios into account, the choice between the `“us”` and `“ucss”` depends on the use case. As a rule of thumb, `“us”` is more suited when steady derivatives are desired, whereas the more stable extrapolation of the `“ucss”` splines should be preferred for *at-line* applications.

### 3.3 Growth rate and timeseries analysis

Most cultivations in microbioreactors such as the BioLector are conducted to extract key performance characteristics of the bioprocesses from the *on-line* measurements. One such performance indicator is the specific growth rate *μ*. In applications where unlimited exponential growth is observed, a constant maximum specific growth rate *μ_max_* can be calculated by regression with an exponential function [5]. Many processes however do not fulfill this assumption and require a more detailed analysis with time-variable specific growth rate. Unlimited exponential growth may be terminated by nutrient limitation, or the characteristics of strain and cultivation media may lead to multiple growth phases. For example, overflow metabolism of *E. coli* growth on glucose can lead to an accumulation of acetic acid which is metabolized in a second growth phase. Accordingly, switchpoints in growth rate can indicate limitations, changes in metabolism or regulation.

From temporally highly resolved backscatter observations combined with a detailed biomass/backscatter correlation model, variable specific growth rate can be calculated using our `bletl.growth.fit_mu_t` function. This model describes the data in a generative fashion by first discretizing time into many segments of exponential growth, followed by simulating the biomass curve resulting from a growth rate that drifts over time. For this it assumes an initial biomass concentration *X*_0_ and a vector of growth rates 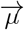, calculates biomass concentrations deterministically and compares them to the observed backscatter using a calibration model built with the `calibr8` package [19]. Parameters *X*_0_ and 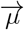 can be obtained through optimization or MCMC. In this analysis we specified a prior belief in *X*_0_ centered around 0.25 g/L, corresponding to typical inoculation density for BioLector experiments. The prior for 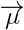 is a random walk of either a Normal or Students-*t* distribution, which pulls the neighboring entries in the growth rate vector closer to each other, resulting in a smooth drift of 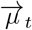 (Section 2.2.2). While this method makes few assumptions about the underlying process and therefore can be applied to many datasets, practitioners wanting to encode process knowledge should also consider differential-equation based modeling approaches for which Python packages such as `pyFOOMB` or `murefi` can be applied [19, 21].

To benchmark the objectivity of the method, we generated a synthetic dataset from a vector of growth rates (Figure 7 **A**, **B**). The comparison of the inference result with the ground truth (Figure 7) shows that with the correct calibration model it yields unbiased estimates of the underlying growth rate. Figure 7 also shows that the `drift_scale` parameter can be tuned to reflect an assumption about the stability of growth rate in the model. Low `drift_scale` constrains the model towards stable exponential growth and correspondingly narrow uncertainties (Figure 7, **C**, **D**). Large `drift_scale` on the other hand encodes the prior belief that growth rate is unstable, leading the model to infer rather unstable growth rates with much higher uncertainty (Figure 7, **E**, **F**).

**Figure 7:**
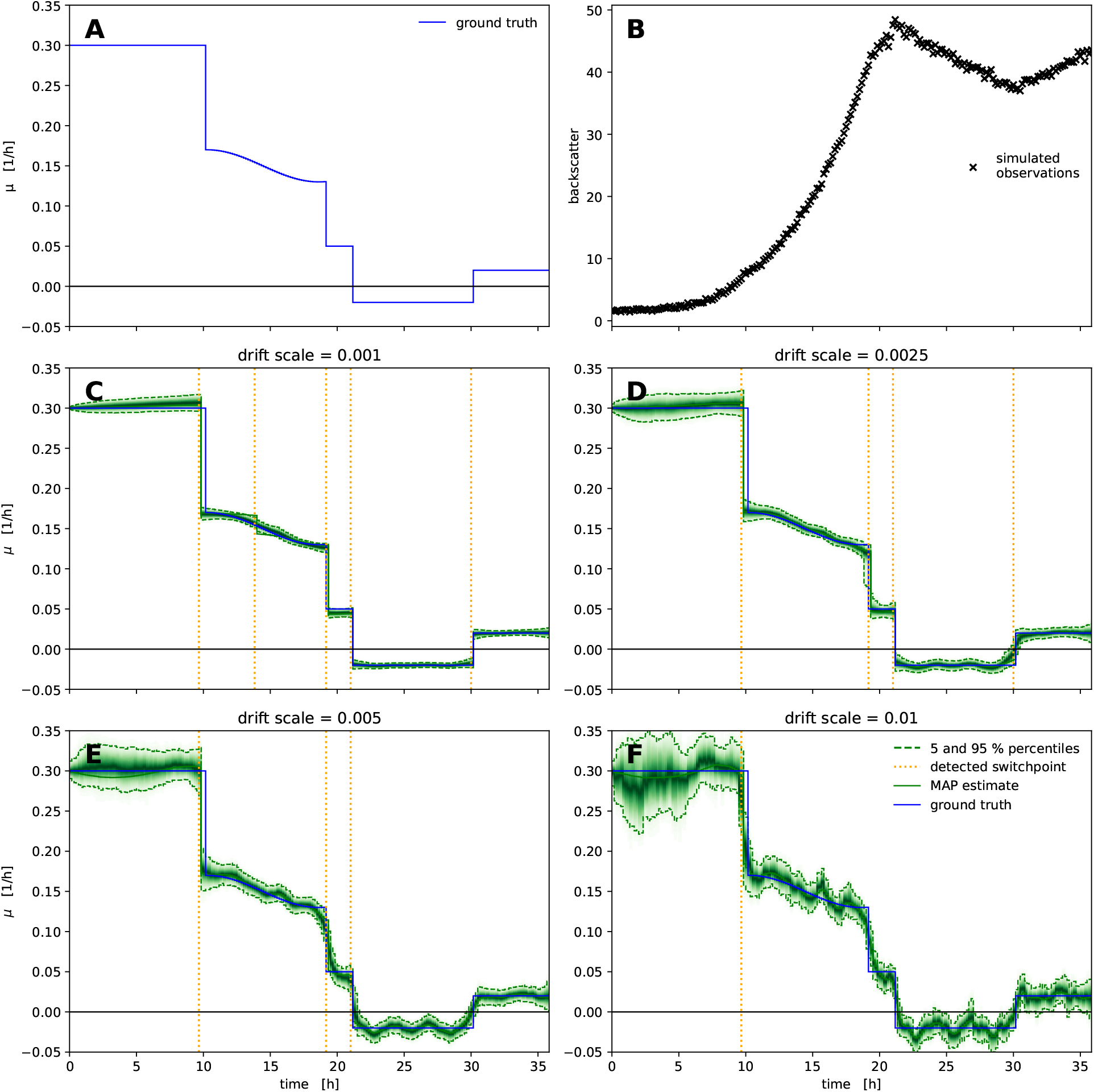
Inference of growth rate from a synthetic dataset. A vector of growth rates (**A**) exhibiting switchpoints and a smooth fluctuation was used to simulate biomass concentrations (not shown) and corresponding backscatter observations (**B**). The magnitude of the `drift_scale` parameter (scale of the Students-t distribution in the random walk) effects stability, switchpoint detection and uncertainty (**C-F**), but in all cases the model fit is unbiased compared to the ground truth. Small `drift_scale` settings constrain the model to stable growth rates, which are inferred with little uncertainty (**C, D**). Large `drift_scale` allows for larger variance in the growth rate, leading to more uncertainty and fewer automatically detected switchpoints (**E, F**). The green density bands visualize the posterior probability density, with dashed lines marking the 5 and 95 % percentiles.

In Figure 8 we applied our generative 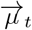 method to data from well F02 of the example dataset. The carbon source composition in this well were 3 parts fructose, 3 parts gluconate and 1 part lactate, causing a change in growth phase at around 9.35 h. The orange line shows the maximum *a-posteriori* estimate of 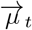, obtained by optimization. Automatically detected growth rate switchpoints are shown as dashed lines. The green density visualizes the percentiles of the posterior probability distribution of the biomass concentration (left) and growth rate (right). The MAP estimate (orange line),is largely in agreement with the full posterior probability distribution obtained by MCMC. This similarity of MAP and the full posterior distribution is not always the case in Bayesian data analysis, but since the computational runtime to obtain the MAP estimate (seconds) is around 100x lower compared to the runtime of a full MCMC parameter estimation (minutes), it is often the first step when analyzing a new dataset.

**Figure 8:**
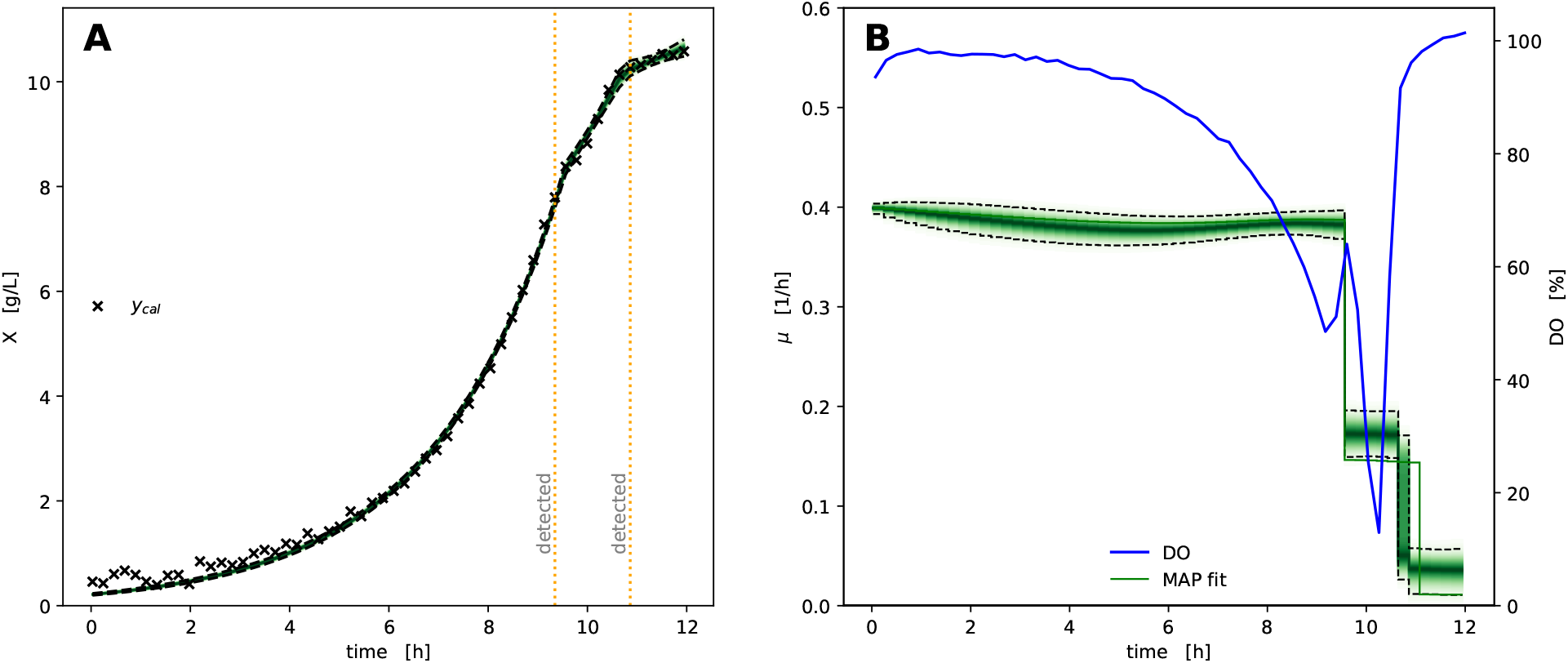
Model prediction of variable growth rate. Biomass concentrations inferred from backscatter observations (**A**) are well explained by the drift of specific growth rate over time (**B**). At two timesteps the specific growth rate changed significantly, which resulted in the automatic detection of switch-points. These switchpoints in 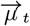 at 9.35 and 10.65 h coincide with changes in the Dissolved Oxygen (DO), indicating a change in cell metabolism. The green density bands visualize the posterior probability density, with dashed lines marking the 5 and 95 % percentiles.

The comparison of growth rate over time (right, orange/green) with dissolved oxygen tension (blue) shows that both detected switchpoints in the growth rate fall together with severe changes in the dissolved oxygen concentration. The first switch from 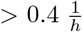 to 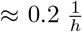 coincides with a temporary increase in DO, whereas the second switch from 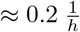 to 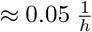 falls together with the final rise in oxygen concentration.

One key aspect of growth rate calculation are the assumptions made about the biomass/backscatter relationship. The aforementioned 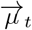 method relies on a *calibration model* of backscatter vs. biomass concentration to simultaneously describe the relationship and measurement noise with a non-linear calibration model. This raises the question to what extend growth rate may be quantified with less sophisticated calibrations.

In Figure 9 we compare the results of a “calibration-free” *μ*(*t*) spline approach (Section 2.2.2) with the 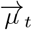 method using linear or logistic calibration models. Note that the “calibration-free” approach also makes the assumption of a linear relationship between biomass concentration and backscatter observations, just without specifying the slope that cancels out in the growth rate calculation (Section 2.2.2).

**Figure 9:**
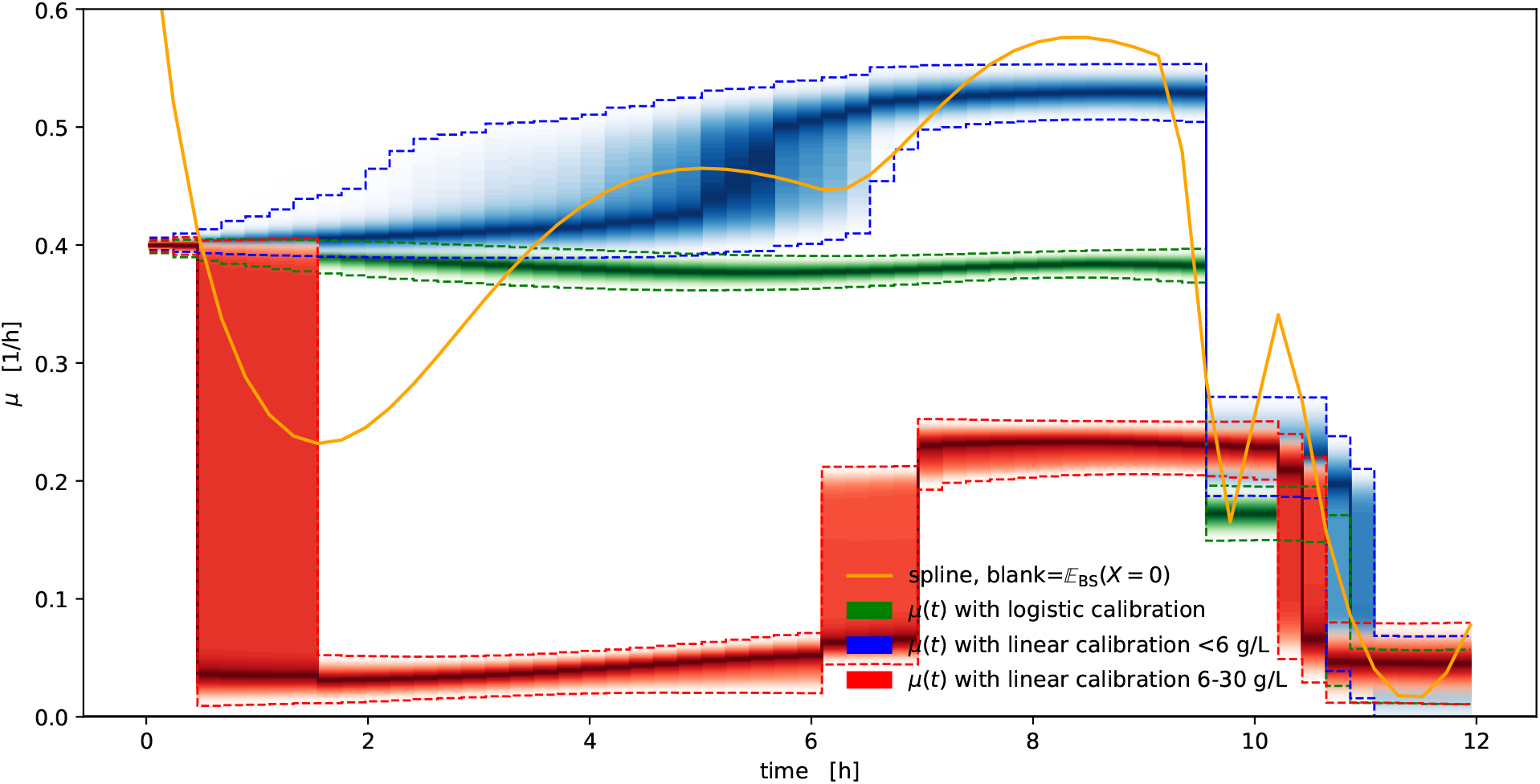
Comparison of growth rate calculation methods. Spline-based *μ*(*t*) growth rate calculation based on blank subtraction (orange) yields a point estimate that fluctuates considerably compared to the “gold standard” of the generative 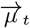 method with detailed biomass/backscatter calibration (green). When the generative method is used with linear calibration models, the choice of calibration concentrations and the decision for (blue) or against (red) fixing the intercept at a blank backscatter has considerable effects on the quality of the outcome. The density bands visualize the posterior probability density, with dashed lines marking the 5 and 95 % percentiles.

Compared to the alternatives, the growth rate curve resulting from the spline method exhibits strong oscillatory artefacts at the beginning of the curve, where the biomass concentration is low. The blue density shows the results of the generative 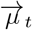 method combined with a linear biomass/backscatter calibration that uses calibration data up to 6 g/L and fixes the intercept to a blank value. This model can still detect the switchpoints, but is biased towards considerably higher growth rates (blue). In contrast, a linear calibration with 6-30 g/L that does not fix the intercept parameter to a blank value leads to a strong under-estimation of the growth rate, largely explained by the lack of fit error of the calibration model (Figure S1). For detailed guidance on the construction and diagnosis of calibration models we refer to [19].

The strength of non-linearities in the biomass/backscatter relationship may depend on the BioLector model and device at hand, but one must conclude that a realistic, unbiased biomass/backscatter calibration is indispensable when quantitative estimates of specific growth rates are desired.

### 3.4 Time series feature extraction

It was previously shown that high-resolution timeseries of culture backscatter can be correlated with product measurements through the use of dimension-reduction techniques and regression models [22]. With `bletl.features` we provide an implementation for configurable and automated extraction of large numbers of features from bioprocess timeseries data. These features may be used as the input to a broad spectrum of machine learning pipelines making use of techniques such as dimension reduction, regression, unsupervised visualization or clustering.

To demonstrate how one might use these methods, we applied them to the previously introduced dataset to obtain a visualization of local structures in the high-dimensional data. Initially 2343 time series features were extracted from the full dataset using both biologically motivated, as well as the statistical time series feature extractors and cleaned to a set of 1282 features (Section 2.2.3). For visualization of the dataset we applied t-distributed Stochastic Neighbor Embedding (t-SNE, Section 2.2.4) to find a 2-dimensional embedding that maintains the local structure of the 1282-dimensional data. The result is a 2-dimensional arrangement (Figure 10) such that culture wells with similar *on-line* signals are located in close proximity.

**Figure 10:**
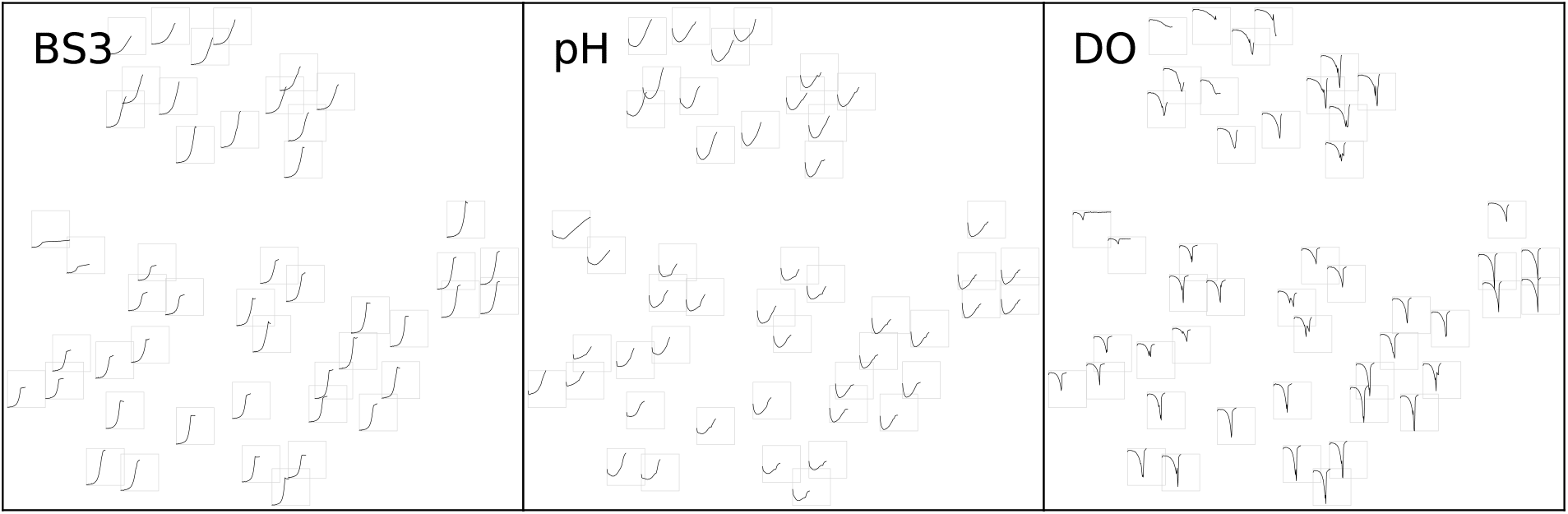
t-SNE from extracted time series features. Each small tile, arranged according to the t-SNE result, shows the time series for one well truncated at the time of harvest. Axis limits of the small tiles are identical within each of the three panels. As with any t-SNE visualization, the large-scale arrangement, rotation or axis units are meaningless, since the technique prioritizes local structure. Note that tiles arranged in close proximity are have similar time series characteristics in all three filtersets.

Coloring the embedding according to the carbon source composition (Figure 11) reveals that the arrangement found by t-SNE is strongly correlated with the presence of gluconate, glutamate and particularly lactate in the cultivation medium.

**Figure 11:**
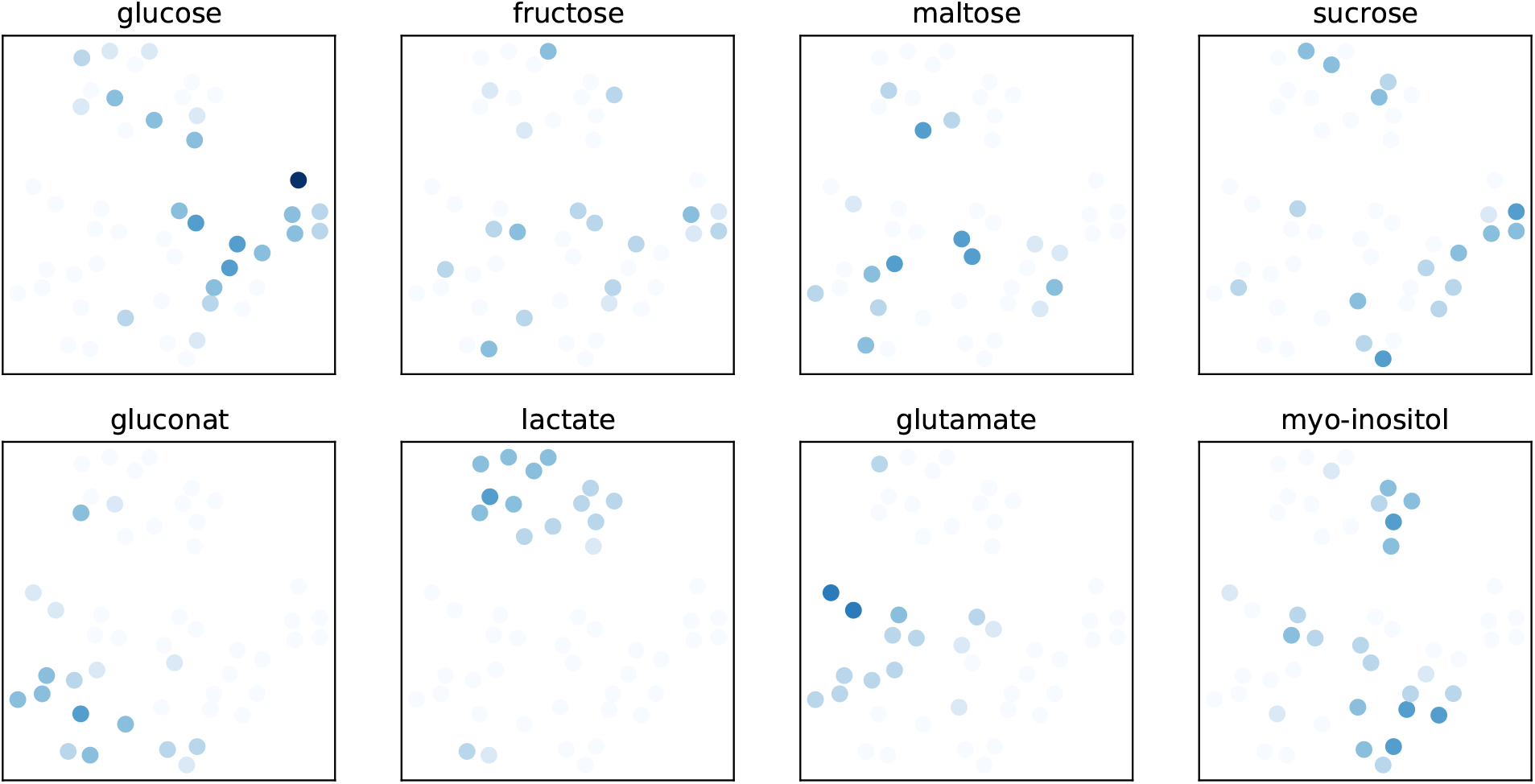
t-SNE embedding colored by carbon sources. Almost all wells that included lactate as a carbon source are closely arranged in the t-SNE embedding, indicating that they are in close proximity in the high dimensional feature space. Likewise, gluconate or glutamate containing wells are closely arranged. In contrast, none of the monomeric or dimeric sugars lead to characteristic BS/pH/DO phenotypes.

The observation that a t-SNE of extracted time series features does not only recover similarities between individual wells, but also aspects of the experiment design shows that our feature extraction is a viable solution to make BioLector datasets amendable to machine learning methods. In contrast to the extraction of manually engineered features [22], our feature extraction workflow works out of the box and with few lines of code.

## 4 Concluding remarks

With the examples in Section 3.1 we showed how `bletl` makes BioLector datasets accessible to standard Python-based data analysis workflows. The switch to Python-based data processing facilitates not only interactive and robust data analysis, but also enables the application of machine learning techniques such as crossvalidated smoothing splines to BioLector datasets. Nevertheless, many scientists who are not yet proficient in Python-based data analysis workflows might be concerned with the initial complexity of the learning curve. That is one of the reasons why the documentation of the `bletl` package comes with ready-to-use examples. The code of the library is thoroughly tested in automated test pipelines to reduce the chance of unexpected failures.

In Section 3.2 we characterized two strategies for smoothing noisy *on-line* signals and showed that subtle differences in implementation can have substantial consequences on the results. This again highlights the need for standardized data structures, robust data analysis routines and thoroughly tested, open-sourced implementations that are distributed through versioned releases.

For the analysis of specific growth rate under not necessarily unlimited exponential growth conditions, we presented a random-walk based 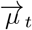 model that can also detect switchpoints automatically. Within seconds our method determines time-variable growth rates by optimization and by leveraging state of the art probabilistic machine learning, it also quantifies Bayesian uncertainties. We showed on a synthetic dataset that the method is not only unbiased, but also offers the practitioner a tuning knob for the bias-variance tradeoff between narrow uncertainties and growth rate flexibility (Figure 7). In comparison with alternative approaches we found that while analyses with less exact, or even without calibration models may still find the same general trends, a quantitative statement about specific growth rate can only be made with accurate calibrations Figure S1.

With Section 3.4 we presented a generally applicable method to extract features for machine learning applications from time series data of microbioreactor experiments. By visualizing the high-dimensional time series features with t-SNE we showed that the features indeed have the information content needed to reconstruct patterns from the experimental design. The visualization of a high-dimensional BioLector dataset in a 2-dimensional arrangement that maintains local structure (Figure 10) is just one example of how our bletl package enriches the exploratory data analysis of microbioreactor experiments.

Overall we conclude that Python packages to parse experimental data into standardized data structures are a valuable asset for quantitative, qualitative and exploratory research.

## Acronyms

bletl: BioLector Extract, Transform, Load.
BS: backscatter.
DBTL: Design – Build – Test – Learn.
DO: Dissolved Oxygen.
FAIR: Findability, Accessibility, Interoperability, Reusability.
IPTG: Isopropyl-*β*-D-thiogalactopyranosid.
MAP: maximum *a-posteriori*.
MBR: microbioreactor.
MCMC: Markov-chain Monte Carlo.
NUTS: No-U-Turn Sampler.
PCA: Principal Component Analysis.
t-SNE: t-distributed Stochastic Neighbor Embedding.

## Acknowledgements

The `bletl` package was developed by Michael Osthege and Niklas Tenhaef. The random-walk 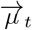 method was devised and implemented by Michael Osthege. Feature extraction and crossvalidation routines were prototyped by Rebecca Zyla using a comprehensive data set provided by Johannes Hemmerich. The strain and experimental workflow for the dataset used in this study were produced by Carolin Müller. Marco Oldiges, Stephan Noack and Wolfgang Wiechert reviewed the manuscript, organized funding and were responsible for supervision and project coordination. This work was funded by the German Federal Ministry of Education and Research (BMBF, Grant. No. 031B0463A) as part of the project “Digitalization In Industrial Biotechnology”, DigInBio. The authors have declared no conflict of interest.

## Appendix

### 4.1 Datasets used in this study

The following dataset files can be found in the data directory of the supporting information GitHub repository [23]:

- `8X4PF4.csv` is the raw BioLector dataset.
- `8X4PF4_eventlog.xlsx` contains metadata of induction and sampling events.
- `8X4PF4_medium_composition.xlsx` is the well-wise composition of carbon sources in the growth media.

### 4.2 Extracted features and t-SNE components

The following files with results from the timeseries feature extraction and embedding are located in the `results` directory of the supporting information GitHub repository [23]:

- `8X4PF4_embedding.xlsx` are the two t-SNE components that were used to arrange data points in the t-SNE visualizations.
- `8X4PF4_extracted_features` contains the raw and cleaned well-wise features extracted from the BioLector dataset.

**Figure S1:**
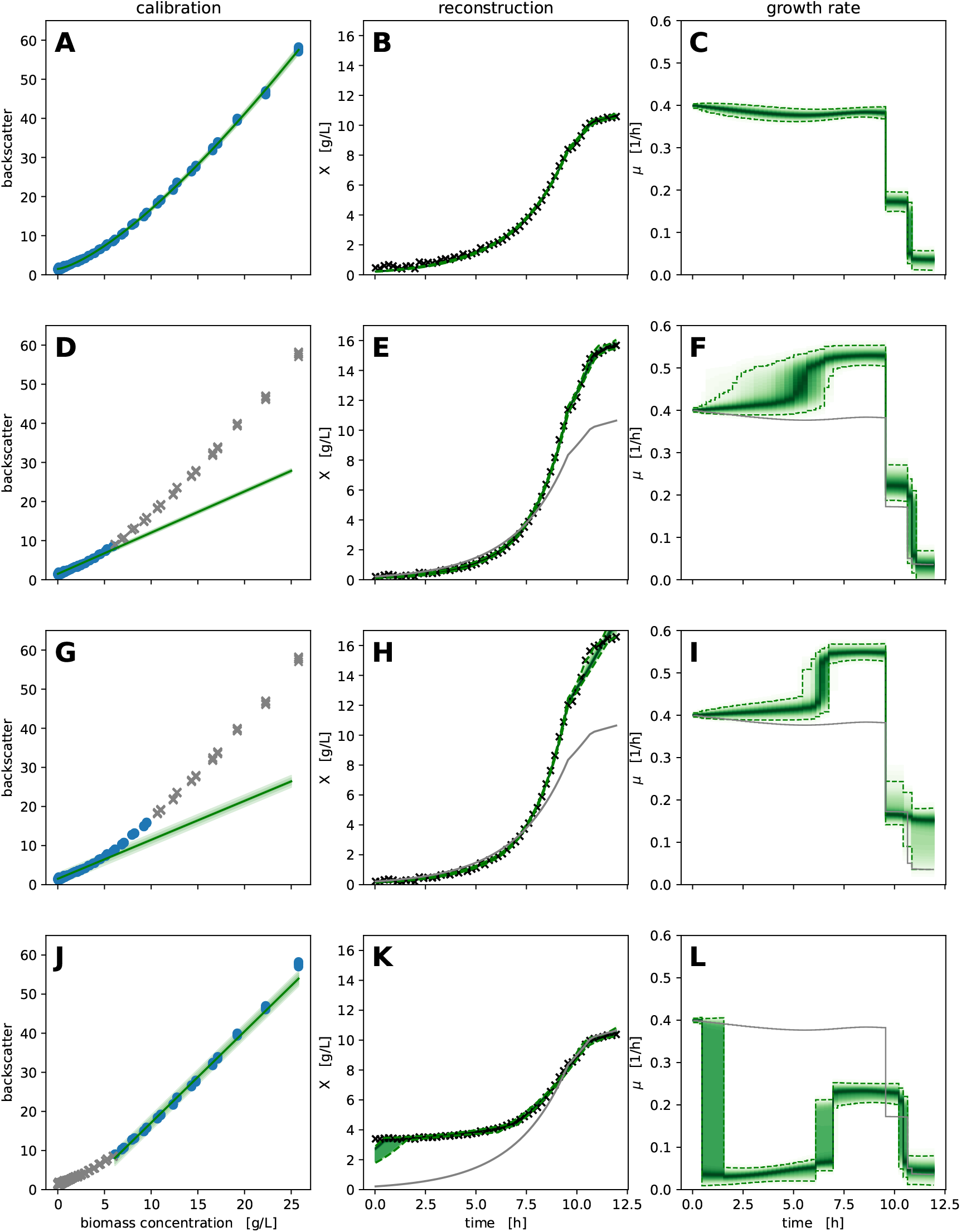
Influence of calibration models on the result of random-walk based 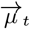 growth rates. An asymmetric logistic calibration model that accurately describes the calibration dataset (**A**) leads to a 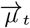 profile (**C**) that is used as the baseline (grey) in the other plots. With a linear calibration model fitted only to calibration data up to 6 g/L (**D**) or the expected maximum biomass concentration of 10 g/L (**G**), the growth rate is systematically overestimated (**F, I**). A linear calibration model through only high biomass concentrations (**J**) that would be amenable to individual cell dry weight determination (e.g. >6 g/L leads to an under-estimation of the growth rate, no longer follows the same profile and does not reliably detect switchpoints (**K, L**). The green density bands of the calibration models (left column) are likelihood bands showing the expected spread of observations, whereas the density bands in the middle and right column show the posterior probability density, with dashed lines marking the 5 and 95 % percentiles.

### 4.3 Plasmid sequence of pCMEx8-NprE-Cutinase

5’-GCACCAATGCTTCTGGCGTCAGGCAGCCATCGGAAGCTGTGGTATGGCTGTGCAGGTCGTAAATCACTGCATAATTCGTGTCGCTCAAGGCGCACTCCCGTTCTGGATAATGTTTTTTGCGCCGACATCATAACGGTTCTGGCAAATATTCTGAAATGAGCTGTTGACAATTAATCATCGGCTCGTATAATGTGTGGAATTGTGAGCGGATAACAATTTCACACAGCAAACAGAATTAAAAGATAGGACCAGGATTACGCCAAGCTTGCAGGCCTGCAGAGAAGACAGGAGAAAAACATATGGGTTTAGGTAAGAAATTGTCTGTTGCTGTCGCTGCTTCGTTTATGAGTTTATCAATCAGCCTGCCAGGTGTTCAGGCTGAATTCCCTACTAGTAACCCTGCTCAGGAGCTTGAGGCGCGCCAGCTTGGTAGAACAACTCGCGACGATCTGATCAACGGCAATAGCGCTTCCTGCGCCGATGTCATCTTCATTTATGCCCGGGGTTCAACAGAGACGGGCAACTTGGGAACTCTCGGTCCTAGCATTGCCTCCAACCTTGAGTCCGCCTTCGGCAAGGACGGTGTCTGGATTCAGGGCGTTGGCGGTGCCTACCGAGCCACTCTTGGAGACAATGCTCTCCCTCGCGGTACCTCTAGCGCCGCAATCAGGGAGATGCTTGGCCTCTTCCAGCAGGCCAACACCAAGTGCCCCGACGCGACTTTGATCGCCGGTGGCTACAGCCAGGGTGCTGCACTTGCGGCCGCCTCCATCGAGGACCTCGACTCGGCCATTCGTGACAAGATCGCCGGAACTGTTCTGTTCGGCTACACCAAGAACCTACAGAACCGTGGCCGAATCCCCAACTACCCTGCCGACAGGACCAAGGTGTTCTGCAATACAGGAGATCTCGTTTGTACTGGTAGCTTGATCGTTGCTGCACCTCACTTGGCTTATGGTCCTGATGCTCGGGGCCCTGCCCCTGAGTTCCTCATCGAGAAGGTTCGGGCTGTCCGTGGTTCTGCTCTTATCGGCTCTGATGGCGGCTCTGGCGGCGGCTCTACATCTCGTGATCACATGGTTCTTCATGAATACGTTAACGCTGCTGGCATCACATAAGAGCTCAATACACTGGCCGTCTTCGTTTTACAGCCAAGCTTGGCTGTTTTGGCGGATGAGAGAAGATTTTCAGCCTGATACAGATTAAATCAGAACGCAGAAGCGGTCTGATAAAACAGAATTTGCCTGGCGGCAGTAGCGCGGTGGTCCCACCTGACCCCATGCCGAACTCAGAAGTGAAACGCCGTAGCGCCGATGGTAGTGTGGGGTCACCCCATGCGAGAGTAGGGAACTGCCAGGCATCAAATAAAACGAAAGGCTCAGTCGAAAGACTGGGCCTTTCGTTTTATCTGTTGTTTGTCGGTGAACGCTCTCCTGAGTAGGACAAATCCGCCGGGAGCGGATTTGAACGTTGCGAAGCAACGGCCCGGAGGGTGGCGGGCAGGACGCCCGCCATAAACTGCCAGGCATCAAATTAAGCAGAAGGCCATCCTGACGGATGGCCTTTTTGCGTTTCTACAAACTCTTTTGTTTATTTTTCTAAATACATTCAAATATGTATCCGCTCATGAGACAATAACCCTGATAAATGCTTCAATAATATTGAAAAAGGAAGAGTATGAGTATTCAACATTTCCGTGTCGCCCTTATTCCCTTTTTTGCGGCATTTTGCCTTCCTGTTTTTGCTCACCCAGAAACGCTGGTGAAAGTAAAAGATGCTGAAGATCAGTTGGGTGCACGAGTGGGTTACATCGAACTGGATCTCAACAGCGGTAAGATCCTTGAGAGTTTTCGCCCCGAAGAACGTTTTCCAATGATGAGCACTTTTAAAGTTCTGCTATGTGGCGCGGTATTATCCCGTGTTGACGCCGGGCAAGAGCAACTCGGTCGCCGCATACACTATTCTCAGAATGACTTGGTTGAGTAATTCGTAATCATGTCATAGGCTCTTCCGCTTCCTCGCTCACTGACTCGCTGCGCTCGGTCGTTCGGCTGCGGCGAGCGGTATCAGCTCACTCAAAGGCGGTAATACGGTTATCCACAGAATCAGGGGATAACGCAGGAAAGAACATGTGAGCAAAAGGCCAGCAAAAGGCCAGGAACCGTAAAAAGGCCGCGTTGCTGGCGTTTTTCCATAGGCTCCGCCCCCCTGACGAGCATCACAAAAATCGACGCTCAAGTCAGAGGTGGCGAAACCCGACAGGACTATAAAGATACCAGGCGTTTCCCCCTGGAAGCTCCCTCGTGCGCTCTCCTGTTCCGACCCTGCCGCTTACCGGATACCTGTCCGCCTTTCTCCCTTCGGGAAGCGTGGCGCTTTCTCATAGCTCACGCTGTAGGTATCTCAGTTCGGTGTAGGTCGTTCGCTCCAAGCTGGGCTGTGTGCACGAACCCCCCGTTCAGCCCGACCGCTGCGCCTTATCCGGTAACTATCGTCTTGAGTCCAACCCGGTAAGACACGACTTATCGCCACTGGCAGCAGCCACTGGTAACAGGATTAGCAGAGCGAGGTATGTAGGCGGTGCTACAGAGTTCTTGAAGTGGTGGCCTAACTACGGCTACACTAGAAGAACAGTATTTGGTATCTGCGCTCTGCTGAAGCCAGTTACCTTCGGAAAAAGAGTTGGTAGCTCTTGATCCGGCAAACAAACCACCGCTGGTAGCGGTGGTTTTTTTGTTTGCAAGCAGCAGATTACGCGCAGAAAAAAAGGATCTCAAGAAGATCCTTTGATCTTTTCTACGGGGTCTGACGCTCAGTGGAACGAAAACTCACGTTAAGGGATTTTGGTCATGAGATTATCAAAAAGGATCTTCACCTAGATCCTTTTGGGGGGGGGGGGAAAGCCACGTTGTGTCTCAAAATCTCTGATGTTACATTGCACAAGATAAAAATATATCATCATGAACAATAAAACTGTCTGCTTACATAAACAGTAATACAAGGGGTGTTATGAGCCATATTCAACGGGAAACGTCTTGCTCGAGGCCGCGATTAAATTCCAACATGGATGCTGATTTATATGGGTATAAATGGGCTCGCGATAATGTCGGGCAATCAGGTGCGACAATCTATCGATTGTATGGGAAGCCCGATGCGCCAGAGTTGTTTCTGAAACATGGCAAAGGTAGCGTTGCCAATGATGTTACAGATGAGATGGTCAGACTAAACTGGCTGACGGAATTTATGCCTCTTCCGACCATCAAGCATTTTATCCGTACTCCTGATGATGCATGGTTACTCACCACTGCGATCCCCGGGAAAACAGCATTCCAGGTATTAGAAGAATATCCTGATTCAGGTGAAAATATTGTTGATGCGCTGGCAGTGTTCCTGCGCCGGTTGCATTCGATTCCTGTTTGTAATTGTCCTTTTAACAGCGATCGCGTATTTCGTCTCGCTCAGGCGCAATCACGAATGAATAACGGTTTGGTTGATGCGAGTGATTTTGATGACGAGCGTAATGGCTGGCCTGTTGAACAAGTCTGGAAAGAAATGCATAAGCTTTTGCCATTCTCACCGGATTCAGTCGTCACTCATGGTGATTTCTCACTTGATAACCTTATTTTTGACGAGGGGAAATTAATAGGTTGTATTGATGTTGGACGAGTCGGAATCGCAGACCGATACCAGGATCTTGCCATCCTATGGAACTGCCTCGGTGAGTTTTCTCCTTCATTACAGAAACGGCTTTTTCAAAAATATGGTATTGATAATCCTGATATGAATAAATTGCAGTTTCATTTGATGCTCGATGAGTTTTTCTAATCAGAATTGGTTAATTGGTTGTAACACTGGCAGAGCATTACGCTGACTTGACGGGACGGCGGCTTTGTTGAATAAATCGAACTTTTGCTGAGTTGAAGGATCAGATCACGCATCTTCCCGACAACGCAGACCGTTCCGTGGCAAAGCAAAAGTTCAAAATCACCAACTGGTCCACCTACAACAAAGCTCTCATCAACCGTGGCTCCCTCACTTTCTGGCTGGATGATGGGGCGATTCAGGCCTGGTATGAGTCAGCAACACCTTCTTCACGAGGCAGACCTCAGCGCCCCCCCCCCCCTAGCTTGTCTACGTCTGATGCTTTGAATCGGACGGACTTGCCGATCTTGTATGCGGTGATTTTTCCCTCGTTTGCCCACTTTTTAATGGTGGCCGGGGTGAGAGCTACGCGGGCGGCGACCTGCTGCGCTGTGATCCAATATTCGGGGTCGTTCACTGGTTCCCCTTTCTGATTTCTGGCATAGAAGAACCCCCGTGAACTGTGTGGTTCCGGGGGTTGCTGATTTTTGCGAGACTTCTCGCGCAATTCCCTAGCTTAGGTGAAAACACCATGAAACACTAGGGAAACACCCATGAAACACCCATTAGGGCAGTAGGGCGGCTTCTTCGTCTAGGGCTTGCATTTGGGCGGTGATCTGGTCTTTAGCGTGTGAAAGTGTGTCGTAGGTGGCGTGCTCAATGCACTCGAACGTCACGTCATTTACCGGGTCACGGTGGGCAAAGAGAACTAGTGGGTTAGACATTGTTTTCCTCGTTGTCGGTGGTGGTGAGCTTTTCTAGCCGCTCGGTAAACGCGGCGATCATGAACTCTTGGAGGTTTTCACCGTTCTGCATGCCTGCGCGCTTCATGTCCTCACGTAGTGCCAAAGGAACGCGTGCGGTGACCACGACGGGCTTAGCCTTTGCCTGCGCTTCTAGTGCTTCGATGGTGGCTTGTGCCTGCGCTTGCTGCGCCTGTAGTGCCTGTTGAGCTTCTTGTAGTTGCTGTTCTAGCTGTGCCTTGGTTGCCATGCTTTAAGACTCTAGTAGCTTTCCTGCGATATGTCATGCGCATGCGTAGCAAACATTGTCCTGCAACTCATTCATTATGTGCAGTGCTCCTGTTACTAGTCGTACATACTCATATTTACCTAGTCTGCATGCAGTGCATGCACATGCAGTCATGTCGTGCTAATGTGTAAAACATGTACATGCAGATTGCTGGGGGTGCAGGGGGCGGAGCCACCCTGTCCATGCGGGGTGTGGGGCTTGCCCCGCCGGTACAGACAGTGAGCACCGGGGCACCTAGTCGCGGATACCCCCCCTAGGTATCGGACACGTAACCCTCCCATGTCGATGCAAATCTTTAACATTGAGTACGGGTAAGCTGGCACGCATAGCCAAGCTAGGCGGCCACCAAACACCACTAAAAATTAATAGTTCCTAGACAAGACAAACCCCCGTGCGAGCTACCAACTCATATATGCACGGGGGCCACATAACCCGAAGGGGTTTCAATTGACAACCATAGCACTAGCTAAGACAACGGGCACAACACCCGCACAAACTCGCACTGCGCAACCCCGCACAACATCGGGTCTAGGTAACACTGAAATAGAAGTGAACACCTCTAAGGAACCGCAGGTCAATGAGGGTTCTAAGGTCACTCGCGCTAGGGCGTGGCGTAGGCAAAACGTCATGTACAAGATCACCAATAGTAAGGCTCTGGCGGGGTGCCATAGGTGGCGCAGGGACGAAGCTGTTGCGGTGTCCTGGTCGTCTAACGGTGCTTCGCAGTTTGAGGGTCTGCAAAACTCTCACTCTCGCTGGGGGTCACCTCTGGCTGAATTGGAAGTCATGGGCGAACGCCGCATTGAGCTGGCTATTGCTACTAAGAATCACTTGGCGGCGGGTGGCGCGCTCATGATGTTTGTGGGCACTGTTCGACACAACCGCTCACAGTCATTTGCGCAGGTTGAAGCGGGTATTAAGACTGCGTACTCTTCGATGGTGAAAACATCTCAGTGGAAGAAAGAACGTGCACGGTACGGGGTGGAGCACACCTATAGTGACTATGAGGTCACAGACTCTTGGGCGAACGGTTGGCACTTGCACCGCAACATGCTGTTGTTCTTGGATCGTCCACTGTCTGACGATGAACTCAAGGCGTTTGAGGATTCCATGTTTTCCCGCTGGTCTGCTGGTGTGGTTAAGGCCGGTATGGACGCGCCACTGCGTGAGCACGGGGTCAAACTTGATCAGGTGTCTACCTGGGGTGGAGACGCTGCGAAAATGGCAACCTACCTCGCTAAGGGCATGTCTCAGGAACTGACTGGCTCCGCTACTAAAACCGCGTCTAAGGGGTCGT ACACGCCGTTTCAGATGTTGGATATGTTGGCCGATCAAAGCGACGCCGGCGAGGATATGGACGCTGTTTTGGTGGCTCGGTGGCGTGAGTATGAGGTTGGTTCTAAAAACCTGCGTTCGTCCTGGTCACGTGGGGCTAAGCGTGCTTTGGGCATTGATTACATAGACGCTGATGTACGTCGTGAAATGGAAGAAGAACTGTACAAGCTCGCCGGTCTGGAAGCACCGGAACGGGTCGAATCAACCCGCGTTGCTGTTGCTTTGGTGAAGCCCGATGATTGGAAACTGATTCAGTCTGATTTCGCGGTTAGGCAGTACGTTCTAGATTGCGTGGATAAGGCTAAGGACGTGGCCGCTGCGCAACGTGTCGCTAATGAGGTGCTGGCAAGTCTGGGTGTGGATTCCACCCCGTGCATGATCGTTATGGATGATGTGGACTTGGACGCGGTTCTGCCTACTCATGGGGACGCTACTAAGCGTGATCTGAATGCGGCGGTGTTCGCGGGTAATGAGCAGACTATTCTTCGCACCCACTAAAAGCGGCATAAACCCCGTTCGATATTTTGTGCGATGAATTTATGGTCAATGTCGCGGGGGCAAACTATGATGGGTCTTGTTGTTGACAATGGCTGATTTCATCAGGAATGGAACTGTCATGCTGTTATGTGCCTGGCTCCTAATCAAAGCTGGGGACAATGGGTTGCCCCGTTGATCTGATCTAGTTCGGATTGGCGGGGCTTCACTGTATCTGGGGGTGGCATCGTGAATAGATTGCACACCGTAGTGGGCAGTGTGCACACCATAGTGGCCATGAGCACCACCACCCCCAGGGACGCCGACGGCGCGAAGCTCTGCGCCTGGTGCGGCTCGGAGATCAAGCAATCCGGCGTCGGCCGGAGCCGGGACTACTGCCGCCGCTCCTGCCGCCAGCGGGCGTACGAGGCCCGGCGCCAGCGCGAGGCGATCGTGTCCGCCGTGGCGTCGGCAGTCGCTCGCCGAGATACGTCACGTGACGAAATGCAGCAGCCTTCCATTCCGTCACGTGACGAAACTCGGGCCGCAGGTCAGAGCACGGTTCCGCCCGCTCCGGCCCTGCCGGACCCCCGGCATCCCGCAAGAGGCCCGGCAGTACCGGCATAACCAAGCCTATGCCTACAGCATCCAGGGTGACGGTGCCGAGGATGACGATGAGCGCATTGTTAGATTTCATACACGGTGCCTGACTGCGTTAGCAATTTAACTGTGATAAACTACCGCATTAAAGCTTATCGATGATAAGCTGTCAAACATGGCCTGTCGCTTGCGGTATTCGGAATCTTGCACGCCCTCGCTCACTGCCCGCTTTCCAGTCGGGAAACCTGTCGTGCCAGCTGCATTAATGAATCGGCCAACGCGCGGGGAGAGGCGGTTTGCGTATTGGGCGCCAGGGTGGTTTTTCTTTTCACCAGTGAGACGGGCAACAGCTGATTGCCCTTCACCGCCTGGCCCTGAGAGAGTTGCAGCAAGCGGTCCACGCTGGTTTGCCCCAGCAGGCGAAAATCCTGTTTGATGGTGGTTAACGGCGGGATATAACATGAGCTGTCCTCGGTATCGTCGTATCCCACTACCGAGATATCCGCACCAACGCGCAGCCCGGACTCGGTAATGGCGCGCATTGCGCCCAGCGCCATCTGATCGTTGGCAACCAGCATCGCAGTGGGAACGATGCCCTCATTCAGCATTTGCATGGTTTGTTGAAAACCGGACATGGCACTCCAGTCGCCTTCCCGTTCCGCTATCGGCTGAATTTGATTGCGAGTGAGATATTTATGCCAGCCAGCCAGACGCAGACGCGCCGAGACAGAACTTAATGGGCCCGCTAACAGCGCGATTTGCTGGTGACCCAATGCGACCAGATGCTCCACGCCCAGTCGCGTACCGTCCTCATGGGAGAAAATAATACTGTTGATGGGTGTCTGGTCAGAGACATCAAGAAATAACGCCGGAACATTAGTGCAGGCAGCTTCCACAGCAATGGCATCCTGGTCATCCAGCGGATAGTTAATGATCAGCCCACTGACGCGTTGCGCGAGAAGATTGTGCACCGCCGCTTTACAGGCTTCGACGCCGCTTCGTTCTACCATCGACACCACCACGCTGGCACCCAGTTGATCGGCGCGAGATTTAATCGCCGCGACAATTTGCGACGGCGCGTGCAGGGCCAGACTGGAGGTGGCAACGCCAATCAGCAACGACTGTTTGCCCGCCAGTTGTTGTGCCACGCGGTTGGGAATGTAATTCAGCTCCGCCATCGCCGCTTCCACTTTTTCCCGCGTTTTCGCAGAAACGTGGCTGGCCTGGTTCACCACGCGGGAAACGGTCTGATAAGAGACACCGGCATACTCTGCGACATCGTATAACGTTACTGGTTTCACATTCACCACCCTGAATTGACTCTCTTCCGGGCGCTATCATGCCATACCGCGAAAGGTTTTGCACCATTCGATGGTGTCAACGTAAATGCATGCCGCTTCGCCTTCGCGCGCGAATTGCAAGCTGATCCGGGCTTATCGACTGCACGGT-3’

## Notes

https://github.com/JuBiotech/bletl-paper

### Competing Interest Statement

The authors have declared no competing interest.

